# Secreted *Aeromonas* GlcNAc binding protein GbpA stimulates epithelial cell proliferation in the zebrafish intestine

**DOI:** 10.1101/2022.06.27.497793

**Authors:** Allison V. Banse, Stephanie VanBeuge, T. Jarrod Smith, Savannah L. Logan, Karen Guillemin

## Abstract

In response to microbiota colonization, the intestinal epithelia of many animals exhibit increased rates of cell proliferation. We used gnotobiotic larval zebrafish to identify a secreted factor from the mutualist *Aeromonas veronii* that is sufficient to promote intestinal epithelial cell proliferation. This secreted *A. veronii* protein is a homologue of the *Vibrio cholerae* GlcNAc binding protein GbpA, which was identified as a chitin-binding colonization factor in mice. GbpA was subsequently shown to be a lytic polysaccharide monooxygenase (LPMO) that can degrade recalcitrant chitin. Our phenotypic characterization of *gbpA* deficient *A. veronii* found no alterations in the mutant cells’ biogeography in the zebrafish intestine and only a modest competitive disadvantage in chitin-binding and colonization fitness when competed against the wild type strain. These results argue against the model of GbpA being a secreted adhesin that binds simultaneously to bacterial cells and GlcNAc, and instead suggests that GbpA is part of a bacterial GlcNAc utilization program. We show that the host proliferative response to GbpA occurs in the absence of bacteria upon exposure of germ-free zebrafish to preparations of native GbpA secreted from either *A. veronii* or *V. cholerae* or recombinant *A. veronii* GbpA.

Furthermore, domain 1 of *A. veronii* GbpA, containing the predicted LPMO activity, is sufficient to stimulate intestinal epithelial proliferation. We conjecture that intestinal epithelia upregulate their rates of renewal in response to secreted bacterial GbpA proteins as an adaptive strategy for coexisting with bacteria that can degrade glycan constituents of the protective intestinal lining.

## INTRODUCTION

Host-associated microbes, collectively called the microbiota, are critical to the development and physiological function of their host animals^1,2^. This complex assemblage of microorganisms contributes to host health in ways that range from stimulating host metabolism to promoting immune system maturation^3^. One way in which microbial communities influence host health is by stimulating cell proliferation of the mucosal epithelia on which they reside. This impact of the microbiota is apparent when comparing epithelial cell proliferation rates of animals raised in the presence (conventionally reared, CV) or absence (germ free, GF) of microbes. For example, GF mice have reduced rates of skin epithelial cell renewal^4^. The animal digestive tract houses the most abundant microbial population and correspondingly the intestinal epithelium shows marked increases in epithelial cell proliferation in CV relative to GF animals, as has been reported in young and adult mice^5,6^, larval zebrafish^7–9^, and larval and adult fruit flies^10,11^. However, the mechanisms underlying microbiota-induced intestinal epithelial cell proliferation are incompletely understood^12^.

Previously, we showed that *Aeromonas veronii*, a common member of the zebrafish intestinal microbiota^16^, secretes an unknown factor(s) that is sufficient to promote epithelial proliferation in the developing intestine of GF zebrafish^8^. The gnotobiotic zebrafish model offers the ability to manipulate the presence^13^ and genetics^14^ of resident microbes in the larval zebrafish, which, combined with the optical transparency and sophisticated genetic tools of the zebrafish model, make it a powerful system to identify bacterial factors that influence aspects of animal tissue development and homeostasis. Our group has used the gnotobiotic zebrafish model to discover specific secreted bacterial proteins that modulate the abundance of intestinal neutrophils^15^ and the expansion of pancreatic beta-cells^16^. Here, we use gnotobiotic zebrafish to identify a secreted *Aeromonas* factor that stimulates intestinal epithelial proliferation, which we show is a homologue of the *Vibrio cholerae* N-acetylglucosamine-binding protein A (GbpA)^17^.

*V. cholerae* GbpA was discovered in a screen for bacterial mutants with impaired adhesion to cultured intestinal epithelial cells^17^ and the *gbpA* mutant was also shown to be defective for binding to chitin-rich zooplankton and chitin-coated beads. Chitin is a complex polymer composed of β-(1→4)-linked N-acetylglucosamine (GlcNAc) monomers. GlcNAs is also a major O-linked glycan component of intestinal mucins, providing a biochemical basis for the parallel binding of *V. cholerae* to chitin- and mucin-rich surfaces. In a neonatal mouse model of infection, the *gbpA* mutant *V. cholerae* were recovered from intestines at lower levels and correspondingly caused less pathology^17,18^. The defective colonization of the *gbpA* mutant was interpreted to be a consequence of its defective adhesion to intestinal epithelia, although such an adhesion defect has not been demonstrated for the mutant during intestinal infection. Unexpectedly for an adhesin, the GbpA protein was found to be a secreted protein^17^. Further structural and biochemical analysis showed that a middle region of the protein (domains 2 and 3) can confer binding of GbpA to *Vibrio* cells^19^, although most of the protein is found in the cell free supernatant. The amino terminal domain of GbpA (domain 1) shares homology with the AA10 family of chitin-degrading lytic polysaccharide monooxygenases (LPMOs)^20^. The C terminal domain (domain 4) resembles a chitin binding domain from *Serratia marcescens* chitinase B and both domains are sufficient to bind to chitin^19^. GbpA’s domain 1 was subsequently shown to be a functional LPMO^21^. The importance of this enzymatic activity of GbpA for *V. cholerae* colonization or pathogenesis has not been explored.

Here we describe the impact of secreted GbpA from *Aeromonas* on intestinal epithelial cells in the context of normal colonization of the microbiota. Using *gbpA* deficient *Aeromonas*, we show that this gene plays no role in bacterial colonization or epithelial adhesion in the zebrafish intestine. We show both *Aeromonas* and *Vibrio* GbpA can stimulate the proliferative response in zebrafish and further demonstrate that the LPMO-containing domain 1 of *Aeromonas* GbpA is sufficient for this activity. Our results demonstrate that the capacity of GbpA to stimulate intestinal epithelial proliferation is intrinsic to this secreted bacteria protein independent of bacterial adhesion or colonization. These finding contribute to a growing appreciation for the role of LPMOs in microbial-host interactions^22^.

## MATERIALS AND METHODS

### Animals

All experiments with zebrafish were performed using protocols approved by the University of Oregon Institutional Animal Care and Use Committee and following standard protocols ^23^. WT (Ab/Tu) zebrafish were reared at 28 °C. GF embryos were derived by surface sterilization of the chorions and maintained as previously described^24^. No exogenous food was provided during the duration of the experiments. CV controls were clutch mates of the GF derived embryos that were not subjected to surface sterilization and were reared in parallel.

### Experimental bacterial strains

To create the *A. veronii ΔgbpA* mutant strain, a vector containing a kanamycin resistance cassette was transformed into SM10 *E. coli*. Conjugation between wild-type *Aeromonas veronii* HM21RS and the vector carrying SM10 *E. coli* strain was carried out, allowing the kanamycin resistance gene to replace the *gbpA* locus in *A. veronii* via allelic exchange.

Candidate mutants were selected for loss of the plasmid and maintenance of kanamycin resistance. Insertion of the kanamycin cassette into the *gbpA* locus was verified in these candidates by PCR. Fluorescently marked derivatives of these strains were engineered with an established Tn7 transposon-based approach^25^. Briefly, a cassette containing the constitutively active synthetic promoter Ptac cloned upstream of genes encoding dTomato or superfolder GFP was chromosomally inserted at the *attTn7* locus to generate *A. veronii attTn7::Ptac-sfGFP* and *A. veronii ΔgbpA attTn7::Ptac-dTomato*. Joerg Graf provided the *A. veronii Δt2ss* mutant and isogenic complementation strain *A. veronii Δt2ss*+*T2SS*^26,27^. Ron Taylor provided *V. cholerae* classical O1 isolate CG842 (strain ATCC 39315 / El Tor Inaba N16961) and isogenic mutant *V. cholerae ΔgbpA* and complementation strain *V. cholerae ΔgbpA+pGbpA*^17^.

### GbpA expression constructs and protein purification

The UniProt ID for GbpA from *A. veronii* strain HM21 is: A0A7Z3TUS8 and the UniProt ID for GbpA from *V. cholerae* strain ATCC 39315 is: Q9KLD5. To generate a plasmid for the expression of unmodified *A. veronii* GbpA, PCR product of the *gbpA* ORF inclusive of the stop codon were generated by using the primers CBPf (gcatcatatggcagcaaaaatccatc)/CBPr1(gcatctcgagtcacttcagctcaatccaggctt). The PCR product was cloned into the NdeI and XhoI sites of plasmid pET21b (Novagen). To generate a plasmid for the expression of a cleavable GST tagged GbpA, PCR product of the *gbpA* ORF lacking the secretion signal and inclusive of the stop codon were generated by using the primers GSTCBPf (gcatgaattccacggctacatcagccagccc)/ GSTCBPr (gcatctcgagtcatcacttcagctcaatccagg) and cloned into the EcoR1 and Xho1 sites of pGEX6p1. The plasmids were then transformed into *E. coli* BL21(DE3) RIL-CodonPlus cells (Stratagene).

Protein purification was achieved using a glutathione Sepharose 4B column (GE Healthcare) following the recommended protocol. Glutathione *S*-transferase (GST) cleavage was achieved after elution of GbpA from the column by adding 1 unit of PreScission protease enzyme and incubation overnight at 4°C per the instructions of the manufacturer (GE Healthcare).

### CFS preparation

Cultures of *A. veronii* strain HM21^27^ in tryptic soy broth (TSB) and *V. cholerae*^17^ in Luria broth (LB) with 0.02% arabinose, pH 6.5, were grown at 30 °C for 17 h on a rotary shaker at 170 rpm. Overnight cultures of *E. coli* BL21(DE3) were grown at 37°C in LB supplemented with 100 μg/ml ampicillin for plasmid maintenance, diluted 1:50 into 50 ml fresh LB/ampicillin, and grown at 37°C until OD_600_ reached ~0.5. IPTG was then added to a final concentration of 1 mM to induce expression of GbpA. The culture was grown with IPTG for 2-3 hours at 30°C. This resulted in a CFS dominated by GbpA, as confirmed via sodium dodecyl-sulfate polyacrylamide gel electrophoresis (SDS-PAGE) followed by Coomassie Brilliant Blue staining, which revealed a dark band of the expected size for GbpA. This band was absent from BL21 CFS carrying an empty pET-21b vector.

Cultures prepared as above were spun at 5,600 × g for 10 min at 4 °C to pellet cells, and the supernatant was passed through a 0.22-μM filter (Corning) on ice. CFS was concentrated through an Amicon Ultra-15 spin concentrator to remove small products, which were toxic to the zebrafish larvae. Protein concentration was determined by Bradford assay. All CFS exposures were performed using ~500 ng/mL total protein.

### *C*hitin binding of *Aeromonas* cells and GbpA protein

To assay *Aeromonas* binding to chitin, magnetic chitin resin (NEB #E8036S) was prepared by washing three times in PBS. Input bacterial suspensions of WT *A. veronii* or *A. veronii ΔgbpA dTomato* (10^9^ colony forming units (CFU)/mL) were applied to the resin and incubated for 30 min or 1 hr at 30°C with gentle rotation. The resin was washed three times with PBS to remove unbound bacteria and finally resuspended in PBS supplemented with 0.4% GlcNAc (Chem-Impex #01427) to release bacteria attached to the resin. Samples were plated on tryptic soy agar (TSA) to calculate the output CFUs and the percent of bacteria attached calculated ([CFUs^output^/CFUs^input^]*100). The competitive binding assay was conducted similarly except equal amounts of WT *A. veronii* and *A. veronii ΔgbpA dTomato* were mixed prior to chitin resin exposure and the CFUs were determined for each strain by visualizing the dTomato expressing colonies using a fluorescent microscope.

To assay GbpA protein binding to chitin, cell free supernatants (CFS) were collected from *E. coli* BL21(DE3) or *V. cholera* following induction of *gbpA* expression as outlined above in CFS preparation with *E. coli* carrying the empty vector (pET-21b) serving as a control. Chitin binding was assayed as described previously^17^. Briefly, normalized CFS was applied to chitin resin and incubated for 1 hr. and the flow through (FT) collected. The resin was washed 5X in PBS, resuspended in 2X protein loading buffer, and boiled for 5 minutes. Proteins in the FT and C fractions were separated via SDS-PAGE and visualized using Coomassie Brilliant Blue.

### Ammonium sulfate fractionation

Ammonium sulfate fractionation was performed on un-concentrated, sterile CFS from 50 mL overnight cultures by slowly adding 100% ammonium sulfate until desired concentration was achieved. These solutions were prepared at 4°C. Precipitated proteins were collected from the 30-40%, 40-50%, 50-60% and 60-70% ammonium sulfate fractions. Precipitated proteins were collected from each fraction by centrifugation at 4°C and 14,000 g for 15 min. The proteins were resuspended in cold embryo medium (EM) and dialyzed for 2–3 hr at 4°C before adding them to 6 days post fertilization (dpf) GF larvae at a final concentration of 500 ng/mL. Pro-proliferative activity was observed in the 50-60% fraction. Hemolysis was assessed by spotting the fractions on blood agar plates.

### Labeling and quantification of proliferating cells

Larvae were immersed in 100 μg/mL 5-ethynyl-2′-deoxyuridine (EdU) solution (A10044; Invitrogen) for 16 h before termination of the experiment. Larvae were fixed in 4% paraformaldehyde for 24 hr at 4 °C, processed for paraffin embedding, and cut into 7-μm sections. For EdU detection, slides were processed according to the Click-iT EdU Cell Proliferation Assay Kit (C35002; Molecular Probes). Samples were imaged on a Nikon Eclipse TE 2000-V inverted microscope equipped with a Photometrics Coolsnap camera. EdU-labeled nuclei within the intestinal epithelium were counted over 30 serial 7-μm sections beginning at the esophageal-intestinal junction and proceeding caudally into the bulb. Analysis of this extended region was necessary because of the stochastic patterns of cell proliferation. The absolute numbers of labeled cells varied between trials. Despite these differences in the absolute numbers of labeled cells, the proportional trends of proliferating cells between treatments were consistent and reproducible between trials.

### Colonization assay

Bacteria were added to GF flasks at 4 dpf at a final concentration of 10^6^ CFUs/mL and incubated with the larvae for 48 hr at 28°C. Larvae were sacrificed at 6 dpf, immediately before the gut was removed and homogenized in a small sample of sterile EM. Dilutions of this gut slurry were plated onto tryptic soy agar and allowed to incubate overnight at 30°C. Colonies from each gut were quantified. A minimum of 10 guts per mono-association or di-association were analyzed.

### Light sheet microscopy

Gnotobiotic larval zebrafish were prepared for light sheet imaging as described by Jemielita et al^28^. Briefly, larvae are anesthetized in dishes filled with sterile EM and tricaine methanesulfonate (MS-222) at 120μg/ml. They were then moved to melted agarose gel and pulled into glass capillaries where the gel was allowed to cool. The capillaries were mounted on a sample holder and a plug of gel containing each live larval zebrafish was extruded into a sample chamber filled with sterile EM and MS-222. Fluid in the sample chamber was maintained at 28°C.

Light sheet fluorescence microscopy was carried out using a home-built light sheet microscope based on the design of Keller et al^29^. Two Coherent sapphire lasers (488nm and 561nm) were rapidly scanned using a galvanometer to create a thin sheet at the focus of an imaging objective. This thin sheet was used to excite fluorescence in specimens. The excitation light was captured as the sample was moved through the sheet, creating a three-dimensional image. To image the entire larval zebrafish gut, four sub-regions were imaged and subsequently registered. A complete image of the gut can be captured in two minutes in two colors with single-micron spacing between planes. All exposure times were 30ms and laser power in both colors was set to 5mW.

### Statistical analysis

Statistical analysis was performed in Prism 9. For comparison of two treatment groups, t-tests were performed. For comparisons across multiple treatment groups, one- and two-way ANOVA analyses were performed as appropriate.

## RESULTS

### *A. veronii* requires the Type II Secretion System for production of a secreted factor that induces intestinal epithelial proliferation

Earlier work from our group demonstrated that *A. veronii* strain HM21 produced an unknown secreted factor(s) that was sufficient to promote cell proliferation in 8 dpf GF larvae, as measured by the number of cells labeled during a 16 hour period of exposure to the nucleotide analogue EdU within a defined 210 μm region of the anterior intestine immediately caudal to the esophageal-intestinal junction^7^ (Figure 1A, B). This factor(s) was present in cell free supernatant (CFS) from *A. veronii* HM21 grown overnight in TSB and fractionated through a spin column to remove small molecular weight material, suggesting the factor(s) was unlikely to be a metabolite and could be a secreted protein^7^. Many Gram-negative bacteria employ the Type II Secretion System (T2SS) to secrete biologically active proteins into the extracellular environment. To test whether the pro-proliferative factor(s) were substrates of the T2SS, we added CFS from an *A. veronii* mutant lacking a functional T2SS (*ΔT2SS*)^26^ or the complement of this mutant with restored T2SS function (*ΔT2SS* + *T2SS*)^26^ to the aquatic environment of 6 dpf GF larval zebrafish and assayed cell proliferation at 8 dpf. We observed that while the CFS from *A. veronii* with a functional T2SS promoted cell proliferation in GF fish similarly to fish with a conventional microbiota, the CFS from the mutant strain lacking T2SS activity was unable to promote cell proliferation above GF levels (Figure 1C). This observation suggests that the pro-proliferative factor(s) is a protein secreted by the T2SS.

**Figure 1.**
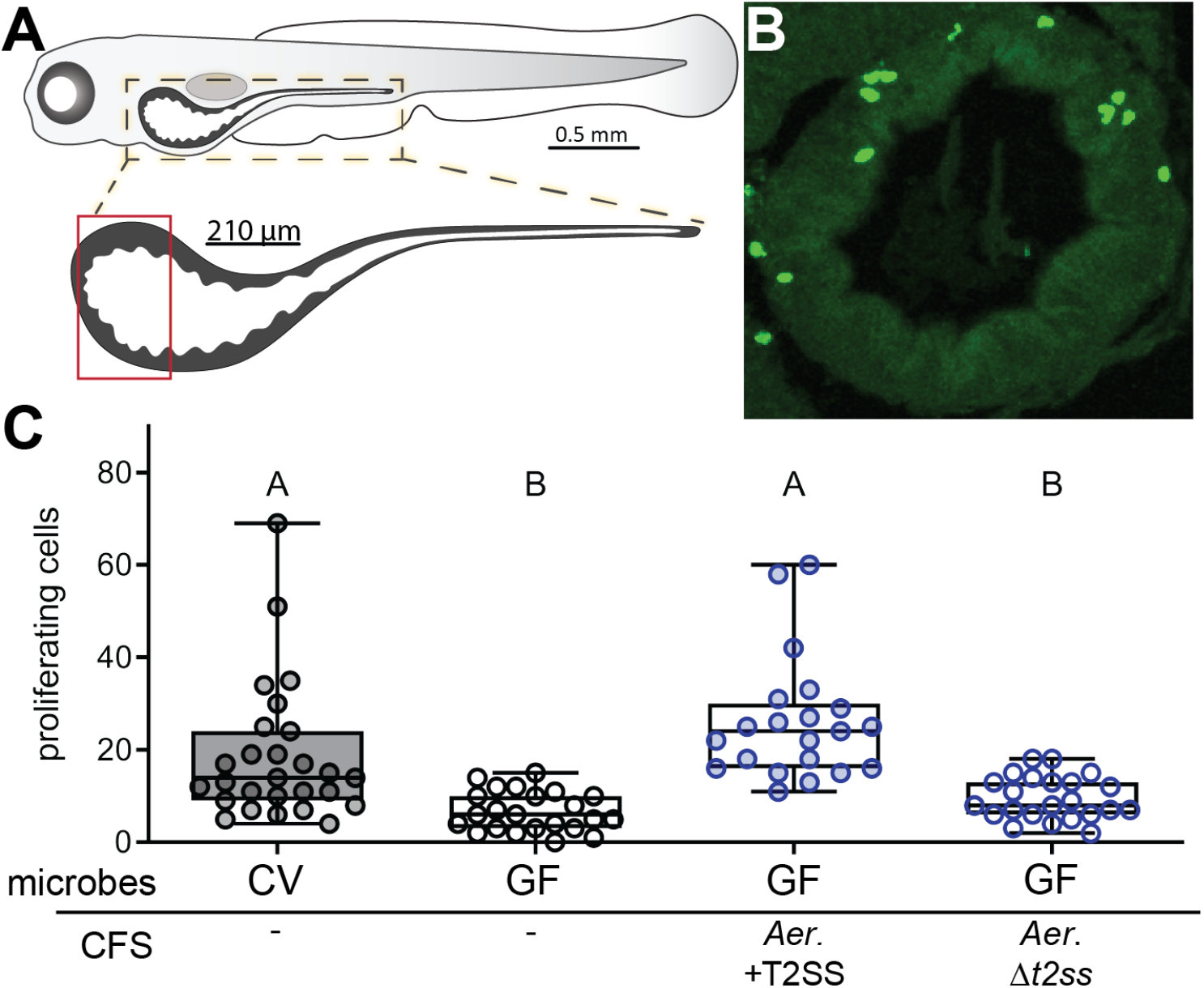
The T2SS of *A. veronii* is required for secretion of a pro-proliferative factor that stimulates intestinal epithelial cell proliferation. **A.** Schematic of the larval zebrafish intestine, highlighting the proximal 210 μm region in which proliferative epithelial cells were quantified, as marked by incorporation of the nucleotide analog EdU. **B.** Representative transverse section of the proximal zebrafish intestine stained to reveal cells that incorporated EdU. **C.** Quantification of proximal intestinal epithelial cell proliferation in 8 dpf CV larvae and 8 dpf GF larvae untreated or exposed from 6 dpf to CFS from *A. veronii* with a functional or mutated T2SS.

### The *A. veronii* pro-proliferative factor is encoded by *gbpA*

To identify candidate pro-proliferative proteins secreted by *A. veronii*, we analyzed a mass spectrometry dataset we had generated of the abundant proteins in the CFS from the T2SS mutant and the complementation strains^15^, focusing on proteins present in the complementation strain and absent in the mutant strain. To further narrow the number of candidate proteins, we fractionated the CFS using ammonium sulfate precipitation, tested these fractions for pro-proliferative activity, and analyzed their composition on Coomassie stained protein gels. We found that the pro-proliferative activity was concentrated in a fraction that appeared to contain a single dominant protein species of approximately 55 kd (Figure 2A). Only two of our candidate proteins identified by mass spectrometry were close to this molecular weight: a hemolysin and a homologue of the *V. cholerae* GbpA secreted protein^17^. We determined that hemolytic activity, assayed on blood agar plates, was concentrated in a fraction lacking pro-proliferative activity (Figure 2A). We therefore turned our attention to GbpA.

**Figure 2.**
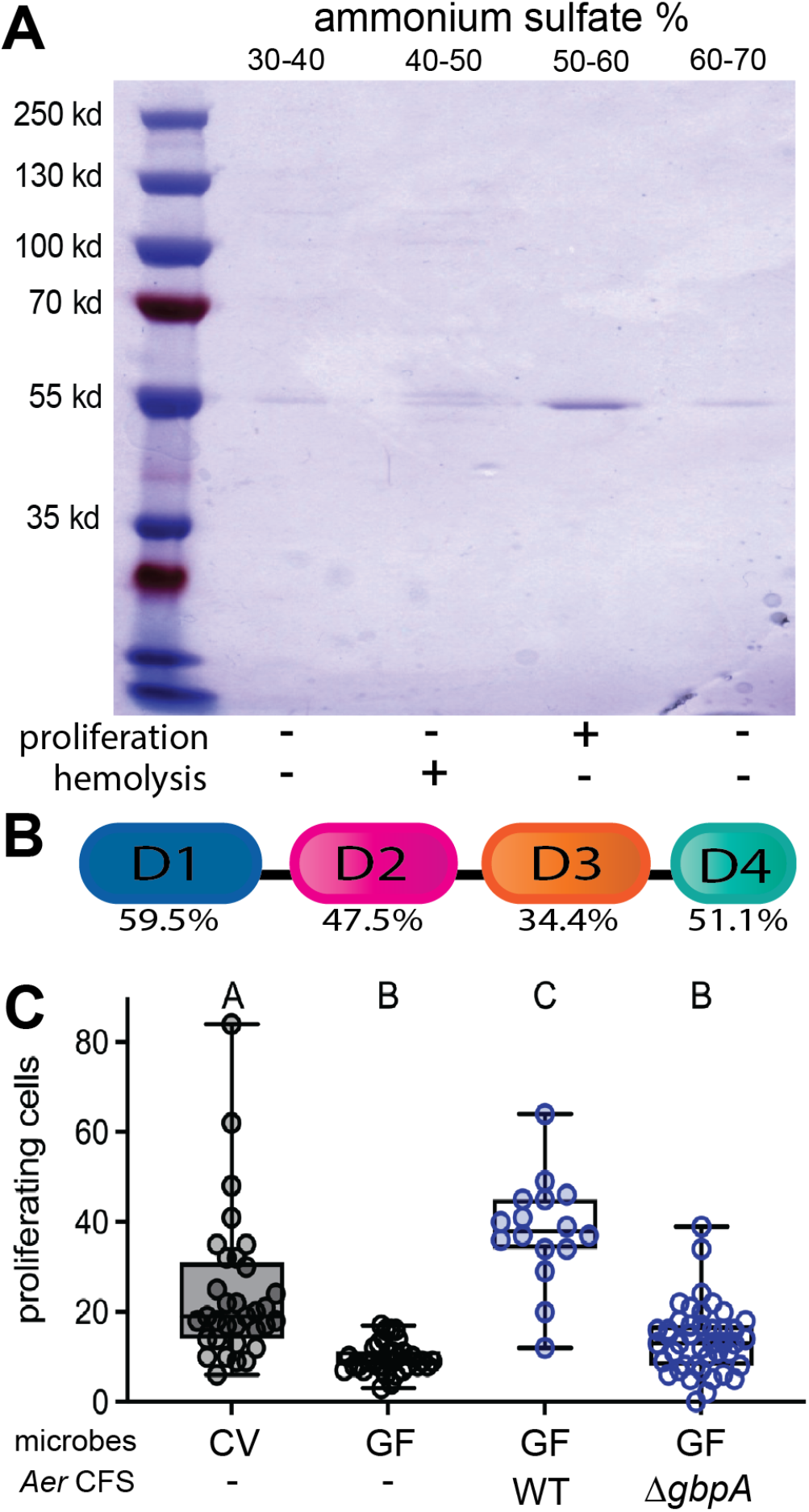
*A. veronii gbpA* encodes the secreted pro-proliferative factor that stimulates intestinal epithelial cell proliferation. **A.** The major protein constituents of ammonium sulfate fractions of *A. veronii* CFS separated by SDS-PAGE and visualized with Coomassie Briliiant Blue, with the corresponding proliferation and hemolysis activity of each fraction indicated below. **B.** Schematic of the shared domain architecture of *A. veronii* and *V. cholerae* GbpA proteins, with the amino acid identity indicated for each of the four protein domains. **C.** Quantification of proximal intestinal epithelial cell proliferation in 8 dpf CV larvae and 8 dpf GF larvae untreated or exposed from 6 dpf to CFS from WT or *ΔgbpA A. veronii*.

GbpA from *A. veronii* has a similar predicted domain architecture to *V. cholerae* GbpA, with low identity putative cell surface binding domains [domain 2 (aa 201-300) and 3 (aa 309-403)] sandwiched between higher identity N-terminal LPMO domain [domain 1 (aa 25-188)] and C-terminal carbohydrate-binding module (CBM) domain [domain 4 (aa 427-468)]. (Figure 2B). The LPMO and CBM domains share 60% and 51% sequence similarity respectively, while domain 3 and 4 are 48% and 34% similar. The LPMO domains of both *Vibrio* and *Aeromonas* GbpA contain two copper-coordinating histidine residues that are necessary for the oxidation reaction which characterizes LPMOs.

To test whether GbpA was necessary for the pro-proliferative activity in the CFS from *A. veronii*, we generated an isogenetic strain of HM21 *A. veronii* in which the *gbpA* open reading frame was replaced by a kanamycin resistance cassette (*gbpA::kan, ΔgbpA*). CFS was collected from the WT and *ΔgbpA* strains and added to the aquatic environment of 6 dpf GF larvae. Whereas the WT CFS elicited a robust proliferative response, the CFS from the *ΔgbpA* mutant failed to induce proliferation above the level observed in GF larvae (Figure 2C), demonstrating that *gbpA* is required for the proliferative response elicited by *A. veronii* CFS.

### Colonization of the zebrafish intestine by *Aeromonas* or *Vibrio* does not require *gbpA*

The *V. cholerae gbpA* mutant was found to exhibit reduced binding to chitin beads following 30 minutes of incubation, as assayed by immunofluorescent microscopy, and reported as number of bacterial cells per bead^17^. We performed a similar chitin bead binding assay with both WT and Δ*gbpA* strains, assessing the fraction of bacteria recovered from chitin beads after both 30 minutes and 1 hour of incubation, using dilution plating. Only about 5% of the population of WT *A. veronii* bound chitin bead at 30 minutes, and this fraction reduced to about 3% by 1 hour (Figure 3A). The Δ*gbpA* population exhibited a lower fraction of chitin bead binding, approximately 0.8%, at both time points (Figure 3A). GbpA was suggested to confer binding of *V. cholerae* cells by a mechanism whereby the protein is first secreted into the extracellular environment, unassociated with the bacterial cell surface, and then subsequently binds to both the bacterial cell and GlcNAc through different protein domains^19,21^. A prediction of this model is that co-incubation of WT and Δ*gbpA* cells strains should rescue chitin-binding defects associated with the Δ*gbpA* mutant, since the predominant form of GbpA from the WT strain is in solution in the culture medium and equivalently accessible to WT or Δ*gbpA* cells. Arguing against the mechanism, we found no change in the percentage of the Δ*gbpA* mutant population that bound to chitin beads when co-incubated with WT cells (Figure 3A). This failure of the WT cells to complement the mutant cells’ chitin-binding defect was also reflected in the competitive indices, calculated as the ratio of the Δ*gbpA* to WT cells recovered from the chitin beads in the mixed incubations, normalized to the ratio of the two strains added initially (Figure 3B).

**Figure 3.**
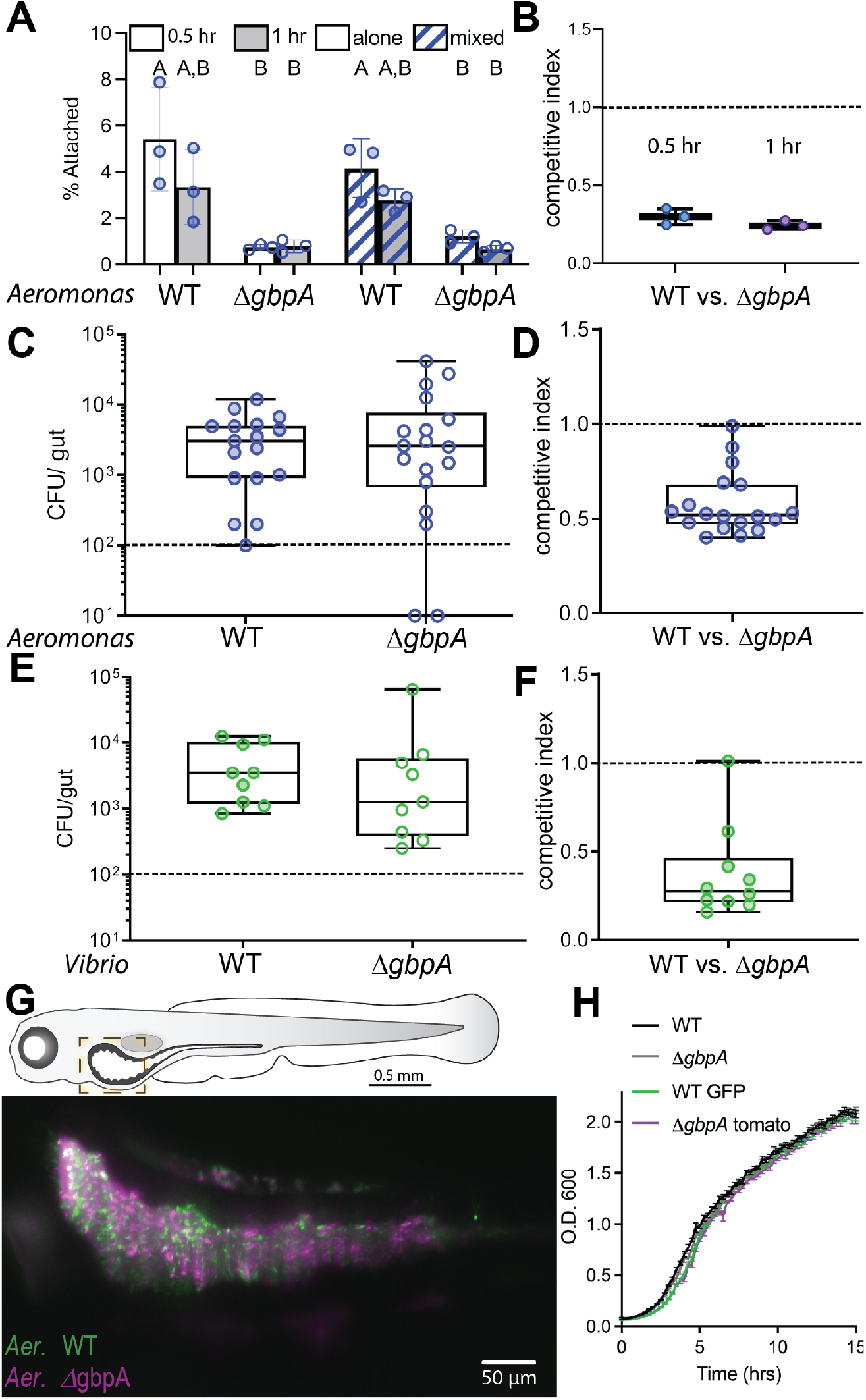
Colonization of the zebrafish intestine by *A. veronii and V. cholerae* does not require *gbpA*. **A.** Binding of WT and *ΔgbpA A. veronii* to chitin beads, quantified as the percent of total bacteria, after either 0.5 hr (white bars) or 1 hour (grey bars) of incubation. Bacterial strains were added to chitin beads alone (solid bars) or mixed with the other strain (striped bars). **B.** Competitive index of *ΔgbpA* versus WT *A. veronii* recovered from chitin beads. **C.** *A. veronii* CFUs recovered at 6 dpf following inoculation of GF zebrafish with individual strains at 4 dpf. **D.** Competitive index of *ΔgbpA* versus WT *A. veronii* recovered at 6 dpf following co-inoculation of GF zebrafish with the two strains at 4 dpf. **E.** *V. cholerae* CFUs recovered at 6 dpf following inoculation of GF zebrafish with individual strains at 4 dpf. **F.** Competitive index of *ΔgbpA* versus WT *V. cholerae* strains recovered at 6 dpf fish following coinoculation of GF zebrafish with the two strains at 4 dpf. **G.** Light sheet micrograph of zebrafish intestine colonized with WT (green) and *ΔgbpA* (purple) *A. veronii* in the proximal intestinal region indicated in the schematic. **H.** Growth curves measuring OD_600_ for each *A. veronii* strain grown in TSB.

In human pathogenic *V. cholerea* strains of both the classic and El Tor biotypes, *gbpA* is required for colonization of the neonatal mouse intestine^17,18^. To test whether *gbpA* was required for *Aeromonas* colonization of the zebrafish intestine, we inoculated the aquatic environment of GF zebrafish on 4dpf, and on 6dpf we assessed the bacterial colony forming units (CFU) per intestine. We observed that both WT and Δ*gbpA* strains were able to colonize the intestine to similar levels, demonstrating that *gbpA* does not play a crucial role in zebrafish gut colonization (Figure 3C).

Certain colonization factors are required only under circumstances of bacterial competition. To test whether *A. veronii gbpA* was required for competitive colonization of the zebrafish intestinal, we co-inoculated WT and Δ*gbpA* strains at equal concentrations to the aquatic environment of GF zebrafish at 4dpf and assayed the CFU/gut of each strain at 6dpf. We calculated the competitive index as the ratio of mutant to WT strains recovered normalized to the ratio inoculated. We observed a modest competitive disadvantage of the Δ*gbpA* mutant when competing against the WT strain (Figure 3D).

*Vibrio* species are normal residents of the zebrafish intestine and human-derived *V. cholerae* can colonize larval zebrafish^30^. We therefore tested a *gbpA* mutant strain of *V. cholerae* that exhibited a mouse colonization defect^17^ in our gnotobiotic zebrafish assay. As with the *A. veronii* strains, we found that both the *V. cholerae ΔgbpA* and WT strains colonized GF zebrafish larvae to similar levels (Figure 3E). When the two *V. cholerae* strains were co-inoculated into GF larvae, we noted a modest competitive disadvantage of the Δ*gbpA* mutant when competing against the WT strain (Figure 3F).

GbpA has been shown to promote the adhesion of *V. cholerae* to cultured intestinal epithelial cells and intestinal tissue explants^17,18^. To test whether *gbpA* was important for *A. veronii* colonization and distribution in the zebrafish intestine, we generated fluorescently labeled strains of both WT (*tn7::GFP*) and the Δ*gbpA* mutant (*gbpA::kan, tn7::dTomato*) and imaged these strains in the intestines of live 6 dpf larval zebrafish using light sheet microscopy^31^. We observed no significant difference in the distribution of the two strains relative to each other or along the zebrafish intestine (Figure 3G). Neither strain exhibited an epithelial proximal distribution but instead were found in bacterial aggregates of a range of sizes within the intestinal lumen, consistent with other imaging we have performed on *Aeromonas* strains in the larval zebrafish intestine^28,31,32^. To exclude the possibility that any of the genetically manipulated *A. veronii* strains had a growth disadvantage under nutrient rich conditions, we compared their growth curves in TSB relative to the WT strain and found that neither the *gbpA::kan* insertion nor the fluorescent protein expression insertions produced any growth defects (Figure 3H). Collectively these results show the Δ*gbpA* mutant has a slight defect in chitin binding and a slight competitive disadvantage in larval zebrafish intestinal colonization, which is not complemented by the presence of GbpA-producing WT cells. Our findings argue against a role for GbpA in *Aeromonas* epithelial adhesion in the larval zebrafish intestine.

### Secreted GbpA from *Aeromonas* and *Vibrio* promote intestinal cell proliferation

Having ruled out a role for GbpA promoting intestinal epithelial cell proliferation through facilitating *Aeromonas* adhesion to the intestinal epithelium, we next explored whether secreted GbpA protein was sufficient to promote intestinal cell proliferation in larval zebrafish. We cloned the *A. veronii gbpA* gene and introduced it on an inducible high copy plasmid into *E. coli*, which lacks any *gbpA* homologues in its genome. Upon induction, a 55 kd protein was the major protein species in the *E. coli* + *pGbpA* CFS, which was absent in the CFS of *E. coli* containing just the empty expression vector. To test whether this recombinant *A. veronii* GbpA had similar chitin binding activity as the *V. cholerea* GbpA protein^17^, we performed chitin binding assays with both proteins. We used CFS from *E. coli* expressing high levels of *A. veronii* GbpA and applied this material to chitin beads. We then collected the flow through (FT) as well as the eluate (E). We observed that *A. veronii* GbpA was present only in the eluate (Figure 4A), indicating that it bound efficiently to the chitin beads. We observed a similar binding to chitin beads when we used CFS from the complementation strain of *V. cholerae ΔgbpA, pGbpA–His* that expresses *V. cholerae* GbpA at high levels from a plasmid pGbpA-His^17^. Consistent with their similar domain architectures, these results indicate the *A. veronii* and *V. cholerae* GbpA proteins share the biochemical property of chitin binding.

**Figure 4.**
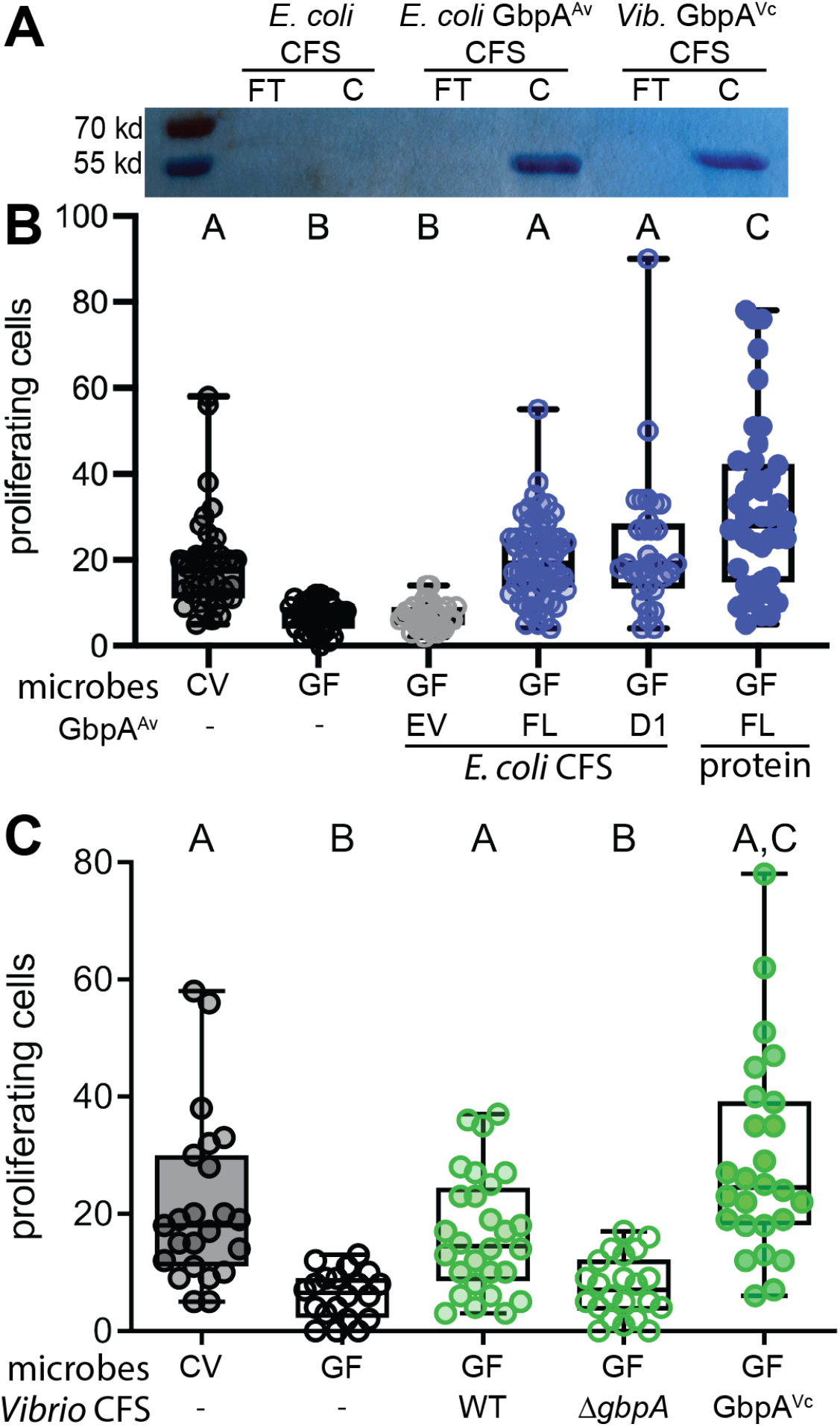
Secreted GbpA proteins are sufficient to increase intestinal epithelial proliferation in GF zebrafish. **A.** CSF from engineered *E. coli* expressing no recombinant protein or IPTG-inducible GbpA^Av^ and from the *V. cholerae gbpA* complementation strain expressing arabinose-inducible GbpA^Vc^ were incubated with chitin beads, rinsed, and the protein content of the flow through (FT) and chitin bead (C) fractions were separated by SDS-PAGE and visualized with Coomassie Brilliant Blue. **B.** Quantification of proximal intestinal epithelial cell proliferation in 8 dpf CV larvae and 8 dpf GF larvae untreated or exposed from 6 dpf to CFS of *E. coli* expressing full length GbpA^Av^, domain 1 of GbpA^Av^, or purified full length GbpA^Av^ protein. **C.** Quantification of proximal intestinal epithelial cell proliferation in 8 dpf CV larvae and 8 dpf GF larvae untreated or exposed from 6 dpf to CFS of WT *V. cholerae*, the *gbpA* deletion strain, and the *gbpA* complementation strain expressing arabinose-inducible GbpA^Vc^.

We next tested the capacity of recombinant *A. veronii* GbpA to induce intestinal epithelial proliferation in GF zebrafish larvae. We collected the CFS from the *E. coli* strains expressing *A. veronii* GbpA or the empty vector and added each to the aquatic environment of 6 dpf GF larvae. We observed that whereas the control CFS had no effect on cell proliferation of GF larvae, the GbpA-enriched CFS promoted cell proliferation to levels like those observed in CV larvae (Figure 4B).

We next explored whether the LPMO-containing domain 1 of *A. veronii* GbpA was sufficient to induce intestinal epithelial cell proliferation. We cloned domain 1 (D1) (aa 25-188) of *A. veronii* GbpA into the inducible expression construct and used a similar strategy to collect CFS from *E. coli* enriched for this protein domain. This CFS containing D1 induced cell proliferation to a similar extent as the full length GbpA from *A. veronii*, indicating that this domain is sufficient to induce the proliferative response in the intestinal epithelium.

To verify that the pro-proliferative activity detected in the *E. coli* CFS was indeed GbpA, we purified a recombinant version of the protein from *E. coli* CFS using a GST tag, which was then removed by proteolysis. When we applied purified GbpA to 6 dpf GF larvae, we observed high levels of cell proliferation at 8 dpf (Figure 4B).

Given the similarity between *A. veronii* and *V. cholerae* GbpA, we next tested whether *V. cholerae* GbpA could also induce a proliferative response in the GF larval zebrafish intestine. We collected CFS from WT, Δ*gbpA* mutant, as well as the complement (*ΔgbpA, pGbpA-His*) *V. cholerae* strains^17^ and applied these each to 6 dpf GF fish. Similar to our observations with *A. veronii*, we observed that the WT and complementation CFS promoted CV-like levels of cell proliferation at 8 dpf, while the mutant CFS did not promote cell proliferation (Figure 4C). This observation suggests that the pro-proliferative activity observed for secreted *A. veronii* GbpA is shared across other GbpA-like proteins produced by resident intestinal bacteria.

## DISCUSSION

Since early descriptions of epithelial cell renewal in the intestines of GF mice^6^, the microbiota has been appreciated as a source of pro-proliferative stimuli that elevates homeostatic rates of intestinal epithelial cell proliferation in many animals. The nature of the microbiota-derived molecules with this activity is incompletely understood. Several bacterial metabolites have been shown to elevate intestinal epithelial proliferations rates in mice and fruit flies, including reactive oxygen species^33^, indoles^34^, and polyamines^35^. Our previous characterization of intestinal epithelial proliferation in gnotobiotic larval zebrafish indicated that *A. veronii*, a prominent bacterial colonizer of the zebrafish intestine, stimulated a proliferative response through secreted factors of greater molecular weight than these metabolites^7^. Here we show that a pro-proliferative factor secreted by *A. veronii* is a homologue of the *V. cholerae* protein GbpA.

The best characterized bacterial proteins that elicit intestinal epithelial cell proliferation are protein toxins of bacterial pathogens. For example, the *Helicobacter pylori* oncogenic virulence factor CagA induces expansion of Lgr5+ stem cells during infection of the gastric epithelium^36^ and transgenic expression of CagA is sufficient to increase epithelial cell proliferation in both zebrafish^37^ and fruit fly^38^ intestines. These pathogen-associated cell proliferative responses, however, are distinct from responses to microbiota colonization in that they are typically associated with inflammation and hypertrophic expansion of the tissue. The proliferative response to the microbiota in the larval zebrafish intestinal epithelium occurs even when tumor necrosis factor (TNF) signaling is blocked^7^, in contrast to the epithelial proliferation in a zebrafish model of spontaneous intestinal inflammation and dysbiosis, which is prevented by interfering with TNF signaling^39^.

GbpA was previously characterized as a virulence factor of human pathogenic *V. cholerae* strains and implicated in disease by a proposed adhesion mechanism of the secreted protein acting to crosslink bacterial cells to GlcNAc moieties on intestinal mucins^17–19^. We generated *gbpA* deficient *A. veronii* to explore the function of this gene in bacterial-host interactions in the larval zebrafish intestine. We found that the *A. veronii ΔgbpA* mutant exhibited reduced binding to chitin beads, as reported for *V. cholerae*, but when we further explored this phenotype in a coincubation assay with our WT and *ΔgbpA A. veronii* strains, we found that this defect was not rescued in trans by GbpA from the WT cells. This unexpected result challenges the current model that GbpA functions primarily as an adhesin since the predominantly soluble GbpA in the extracellular environment should be readily accessed by both the WT and *ΔgbpA* mutant cells. Additional evidence challenging the idea that GbpA’s major function is as a chitin adhesin comes from a previous study showing that a *V. cholerae fliA* mutant upregulates *gbpA* expression but is defective for chitin binding compared to the WT strain due to downregulation of the cell surface associated adhesin FrhA^40,41^. We also found no evidence for GbpA conferring a colonization advantage to either *A. veronii* or *V. cholerae* in the larval zebrafish intestine when introduced in mono-associations, but a slight competitive disadvantage in the presence of GbpA-secreting WT strains. Finally, we found no evidence that GpbA expression altered the intestinal biogeography of *A. veronii*, which normally colonizes the intestinal lumen in cellular aggregates. Our observations argue against the model that *A. veronii* utilizes GpbA as an adhesion to colonize GlcNAc-rich surfaces. Instead, we hypothesize that GbpA is part of a GlcNAc utilization program that *A. veronii* deploys when GlcNAc is an advantageous carbon source. In support of this idea, *V. cholerae* has been shown to upregulate *gbpA* in the presence of GlcNAc and chitin oligosaccharides^42^.

We predict the binding defects observed on chitin resin reflect physiological differences between the WT and *ΔgbpA Aeromonas* due to alterations in nutrient processing and acquisition in the *ΔgbpA* mutant. This idea is consistent with a recent characterization of *Pseudomonas aeruginosa* mutants lacking the gene for Chitin binding protein D (CbpD), a LPMO with similar N and C terminal chitin binding domains as GbpA^43^. The authors showed that *cbpD* deficient *P. aeruginosa* have markedly altered transcriptomes and proteomes as compared with WT cells grown in various media, and the altered metabolic state of the cells correlated with reduced survival in blood and pathogenicity in mouse tissues. This new model for GbpA function helps resolve the longstanding question of how a protein found predominantly in the extracellular environment, unattached to the bacterial cell surface, contributes to *V. cholerae* colonization and pathogenesis.

Independent of possible fitness advantages conferred by GbpA to *A. veronii* during host colonization, we show that the secreted protein stimulates epithelial proliferation in the GF larval intestine. This pro-proliferative activity is recapitulated by the LPMO-containing domain 1 of the protein and by the related *V. cholerae* GbpA. We hypothesize that a consequence of *A. veronii* secreting GbpA in the intestine as part of a GlcNAc utilization program is that the enzymatic activity or byproducts are sense by the host through innate immune pathways that monitor intestinal mucosal integrity or glycan composition. We have shown that the host proliferative response to the microbiota requires the innate immune adaptor Myd88, but that activation of innate immune signaling with bacterial lipopolysaccharide (LPS) is not sufficient to elicit intestinal epithelial cell proliferation^7^. In the zebrafish intestine, GbpA would have access to chitin as a component of the zebrafish intestinal lining^44^. Chitin is a feature of all invertebrate intestines and many vertebrate intestines, where it coexists in varied proportions with a meshwork of glycan-rich mucins^45^. Even in mammalian intestines that lack a chitin layer, the intestinal mucus contains ample GlcNAc polysaccharides that could be cleaved by bacterial secreted LPMOs.

As a bacterial protein that can target the intestinal lining, GbpA represents an example of a Microbial Associated Competitive Activity (MACA), a term we coined to describe microbial activities important for competitive fitness in multi-species communities that are sensed by host tissues as sources of information for regulating programs of development and repair^46^. In this regard, the intestinal epithelial proliferation observed upon microbiota colonization is not an outcome that host-associated bacteria LPMOs evolved to evoke, but rather an adaptation of the host tissue to bolster epithelial renewal programs in the face of degrative enzymes secreted by specific constituents of its intestinal microbiota. The MACA framework explains how specific secreted proteins from microbiota members can have profound impacts on host tissue development and physiology and how different nonhomologous proteins can elicit similar effects by executing similar activities. Future studies of GbpA will test whether it elicits host epithelial proliferative response via its LPMO activity and whether specific byproducts of its enzymatic reaction or depletion of its co-substrates dioxygen or hydrogen peroxide are sensed by the host to induce epithelial renewal programs.

## Author contributions

**AVB**: conceptualization, methodology, investigation, original draft preparation; **SV**: visualization, original draft preparation; **TJS**: investigation (chitin binding assay), manuscript review and editing; **SLL**: investigation (light sheet microscopy), visualization; **KG**: conceptualization, funding acquisition, supervision, original draft preparation, review and editing.

## Funding Sources

Research reported in this publication was supported by the NIH under award numbers F32DK096755 (to AVB), 5T32GM007413 (to SVB), F32DK124033 (to TJS), and 1R01 CA176579, 1P50GM098911, and 1P01GM125576 (to KG). The content is solely the responsibility of the authors and does not necessarily represent the official views of the NIH.

## ACKNOLWEDGEMENTS

We thank Joerg Graf for the generous gift of *A. veronii* strains, Ron Taylor for the generous gift of *V. cholerae* strains, JT Neal for the micrograph of EdU labelled intestine, Erika Mittge for assistance with gnotobiotic zebrafish experiments, W. Zac Stephens for assistance with *A. veronii* genetics, the UO Histology Facility for tissue sectioning and Rose Sockol and the UO Zebrafish Facility for maintenance of zebrafish lines.

## ABBREVIATIONS

GlcNAc: N-acetylglucosamine
GbpA: GlcNAc binding protein A
LPMO: lytic polysaccharide monooxygenase
CFS: cell free supernatant
CV: conventionally reared
GF: germ free
WT: wild type
CFU: colony forming unity
TSB: tryptic soy broth
TSA: tryptic soy agar
LB: Luria broth
EM: embryo medium
MS-222: tricaine methanesulfonate
dpf: days post fertilization
EdU: 5-ethynyl-2′-deoxyuridine
SDS-PAGE: Sodium dodecyl-sulfate polyacrylamide gel electrophoresis
ANOVA: analysis of variance.

## Notes

### Competing Interest Statement

The authors have declared no competing interest.

